# BIOFABRICATION OF AN OVINE INTERVERTEBRAL DISC MODEL BY COMBINING A POLYCAPROLACTONE FRAME WITH A BIOPRINTED ALGINATE HYDROGEL

**DOI:** 10.1101/2025.03.13.642855

**Authors:** Emmaëlle Carrot, Mansoor Chaaban, Daronne Cano Contreras, Clara Schiex, Joëlle Véziers, Boris Halgand, François Loll, Johann Clouet, Michael G. Monaghan, Marion Fusellier, Jérôme Guicheux, Vianney Delplace, Catherine Le Visage

## Abstract

The intervertebral disc (IVD) primarily comprises an outer ring of collagen fibers (annulus fibrosus, AF), which encases a soft, gelatinous core (nucleus pulposus, NP). Existing in vitro models have failed to integrate these two tissues effectively or accurately replicate their intricate organization. By combining two biofabrication techniques, we developed a novel 3D in vitro model that closely mimics the organization of an ovine IVD. Our approach employs a polycaprolactone (PCL) frame produced via melt electrowriting to recreate the multilamellar architecture of the annulus fibrosus. Ovine primary cells, encapsulated in a photocrosslinkable alginate hydrogel, were precisely extruded within the multilamellar structure, thereby mimicking the native shape and size of an ovine disc. The bioink containing the NP cells was deposited at the center of the construct, while the bioink with the AF cells was strategically layered in between the lamellae of the PCL frame. Photocrosslinking was optimized to match the native stiffness of the disc. The constructs were maintained in culture for 28 days, during which we thoroughly assessed reproducibility, stability, and cell viability and phenotype. The results unequivocally demonstrated that the PCL frame effectively guided the alignment and proliferation of AF cells, while the alginate hydrogel preserved NP cell phenotype. This model successfully replicates the organization of the IVD, providing a promising platform for advancing our understanding of disc biology and driving the development of novel therapeutic strategies.

## 1. Introduction

The intervertebral disc (IVD) is a fibrocartilaginous tissue that supports the junction between adjacent vertebrae and enhances spinal flexibility by permitting movements such as flexion, extension, lateral bending, and rotation [1]. The functions of the IVD closely correlate with its specific internal organization, which is divided into two main regions sandwiched between the vertebral endplates. The nucleus pulposus (NP), which constitutes the core of the IVD, is a highly hydrated tissue rich in proteoglycans and randomly arranged type II collagen fibers. The NP is a viscoelastic tissue displaying a compressive stiffness, or Young’s modulus, of around 5 kPa [2]. In the human adult IVD, NP cells are rounded chondrocyte-like cells with a density of 4,000 cells/mm3. These cells maintain the extracellular matrix (ECM) through synthesizing ECM macromolecules, including aggrecan and type II collagen. The NP is surrounded by a highly organized fibrous tissue named annulus fibrosus (AF). This tissue mainly comprises type I collagen fibers aligned and organized in concentric lamellae [3]. Their number varies between 15 and 25, depending on the location in the IVD, the spine region, and age [4]. This anisotropic organization allows the AF to withstand multidirectional loads and maintain the integrity of the disc [5]. The cell density is around 3-fold higher in the AF than in the NP, with an average of 12,000 cells/mm3 [6]. AF cells exhibit an elongated morphology and align parallel to the oriented collagen fibers [7]. Similar to the NP cells, the AF cells are crucial for tissue homeostasis by preserving the composition and organization of the ECM. These two regions facilitate the absorption and distribution of stress in the spine. The endplates separate the IVD from the vertebral bone and prevent the highly hydrated nucleus from bulging into the adjacent vertebrae.

With aging, there is a transition from anabolism to catabolism characterized by an alteration in the cell number and the activation of inflammatory pathways, leading to tissue degradation and loss of structure and function of the IVD [8,9]. IVD degeneration is a leading cause of low back pain [10,11]. The Ovine is mainly used as a preclinical model because of its biomechanical and cellular similarities with the human [12]. Despite the significant social, medical, and economic impact of low back pain, the current conservative and surgical approaches focus on treating the symptoms instead of intervening in disc degeneration and delaying its progression [13]. New therapeutic approaches are being developed to address this issue. These strategies include cells, small molecules, nucleic acids, or biomaterials [14]. None have been approved for clinical use, and most have failed preclinical studies. This is partly due to the lack of relevance of the models used for the evaluation.

Currently, most therapeutic strategies for IVD degeneration are evaluated on 2D cell monolayer systems [15]. These systems provide information on the toxicity and effect of the therapeutic strategy at the cellular level. However, these monolayer systems do not replicate the spatial cellular organization of the native IVD, and cells exhibit phenotypic instabilities due to the absence of an ECM-rich 3D microenvironment, which limits the applicability of results to *in vivo* situations [16]. These drawbacks have prompted researchers to reconsider conventional culture systems, notably to increase their relevance and predictivity. Therefore, 3D cell culture has been proposed as a potential *in vitro* system. Among the different options for culturing cells in a 3D environment, encapsulating the cells in hydrogel-based systems that mimic the ECM is most commonly used. It has already been shown that NP cells encapsulated in alginate beads maintain their specific phenotypic expression and deposit matrix in the beads after several days [17–20]. Other biomaterials have been considered for building 3D cell culture systems, such as fibrin [21], cellulose [22] and collagen [23]. Although these 3D models are physiologically more relevant than 2D culture models, they still need improvement as they still lack key requirements to capitulate IVD characteristics. For example, the size of these models does not represent physiological conditions, their shape does not mimic that of the IVD, and these models generally represent a single cell type which does not recapitulate IVD cellular composition.

In this work, we aim to present an innovative biofabrication strategy of an ovine IVD *in vitro* model that replicates the spatial native organization and phenotype of NP and AF cells. A polycaprolactone (PCL) frame obtained by melt electrowriting (MEW) was designed to mimic the AF lamellar structure and guide cell alignment in the fabricated model. Cells were encapsulated into an alginate-methacrylamide (AlgMA) hydrogel, which achieved tissue stiffness once photocrosslinked. Extrusion-based bioprinting was then used to precisely control the spatial deposition of AF cell-loaded AlgMA within the PCL scaffold and NP cell-loaded AlgMA in the center of the construct to form the IVD construct. The biological behavior of ovine NP and AF cells was explored and compared with native ovine tissues.

## 2. Materials and methods

### 2.1 Materials

2-aminoethyl methacrylamide hydrochloride (900652), 2-(N-morpholino)ethanesulfonic acid (MES, M3671), 2-hydroxy-4’-5ethydroxyethoxy)-2-methylpropiophenone (IRG, 410896), Lithium phenyl-2,4,6-trimethylbenzoylphosphinate (LAP, 900889), 2-phospho-L-ascorbic acid trisodium salt (2AA, 49752), hyaluronidase (H4272), trypsin (T9935), collagenase (C5138), PCL (Mn 80,000, 440744), Triton™ X-100 (T8787), freon (254991) and sucrose (S7903) were purchased from Sigma-Aldrich. 4-(4,6-Dimethoxy-1,3,5-triazin-2-yl)-4-methylmorpholinium tetrafluoroborate (DMT-MM, D2919) were purchased from TCL chemiacals. Sodium alginate (Protanal LF10/60, MW = 60-180 KDa) was purchased from IMCD. Dulbecco’s Phosphate Buffered Saline (DPBS, 21600069), Dulbecco’s Modified Eagle Medium (DMEM)-high glucose, GlutaMAX™, pyruvate (31966047), penicillin/streptomycin (P/S, 10,000 U/mL, 15140122), amphotericin B (250 μg/mL, 15290026), phalloidin-Alexa Fluor™ 647 (A22287), Syto 13™ Green-Fluorescent Nucleic Acid Stains (Syto 13™, S7575), Live/dead assay (L3224), Monomeric Cyanine Nucleic Acid Stains (TO-PROTM-3, T3605), Alginate lyase (A1603), bovine serum albumin (BSA, A9647) and Tween® 20 (P2287) were purchased from ThermoFisher Scientific. Glutaraldehyde (AGR1012) and osmium (AGR1021) were purchased from Agar Scientific Ltd. MTT assay (475989) was purchased from Merk Millipore. Fetal serum bovine (CVFSVF00 01) was purchased from Eurobio. Hanks’ Balanced Salt Solution (HBSS, L0606) was purchased from Biowest. Anti-collagen type I (ab138492), anti-collagen type II (ab34712), and anti-rabbit Alexa Fluor 568 (ab175471) were purchased from Abcam.

### 2.2 Synthesis of alginate-methacrylamide (AlgMA)

Alginate-methacrylamide (AlgMA) was synthesized by chemically modifying sodium alginate with 2 aminoethyl methacrylamide hydrochloride through amidation. Briefly, a 1% (w/v) alginate solution was prepared in MES buffer (pH 5.5) under stirring at room temperature. After complete dissolution, DMT MM (1 equivalent to carboxylic acids) was added and allowed to react for 30 min to activate the carboxylic acid groups. Finally, 2 aminoethyl methacrylamide hydrochloride (0.5 equivalent to carboxylic acids) was added and allowed to react for 3 days at room temperature while stirring. The solution was dialyzed against 0.1 mol/L DPBS for 24 h at room temperature, then against deionized water for 48 h. Finally, the solution was sterilized through a 0.22-µm-pore filter, lyophilized, and stored at - 20 °C. Elemental analysis (ThermoFisher, Flash 2000) was used to quantify the nitrogen: carbon ratio in AlgMA and determine the degree of substitution.

### 2.3 Synthesis and mechanical characterization of AlgMA biomaterial ink

Various photoinitiators (LAP or IRG, 10 mmol/L; AlgMA 2%), energy density (4, 10, 12 or 15 mJ/mm^2^, 405 nm; LAP 10 mmol/L; AlgMA 2%) and AlgMA concentration (1%, 2%, 4% or 5% [w/v]; LAP 10 mmol/L; 405 nm, 10 mJ/mm^2^) were tested. AlgMA was dissolved in the culture medium containing the photoinitiator. Thirty µL of an AlgMA/photoinitiator solution was cast into a mold of 1.5 mm (*h,* height) x 5 mm (*Ø*, diameter). The solution containing IRG was cross-linked with a 365 nm UV lamp (UVP, B-100AP, power: 20 mW/cm^2^) while the one containing LAP was cross-linked with either a 365 nm UV lamp or a 405 nm lamp (RegenHu, 900 019 386, power: 50-100 mW/cm²). A 5-second-increment cross-linking experiment was used to determine the gelation time needed to form AlgMA hydrogels that can be uncasted from the mold with a spatula. The determined gelation time was used for all subsequent tests. Hydrogel stiffness was measured with an unconfined compression test using a MicroTester® (CellScale). The compression measurements were performed under 20% strain. Each hydrogel underwent a 30-second compression cycle using a 6 mm square stainless-steel plate placed upon a 58 mm long circular tungsten microbeam (558.8 μm diameter). The force (μN) and the displacement (μm) were recorded, and Young’s modulus was calculated according to the manufacturer’s recommendations and the standard expression:

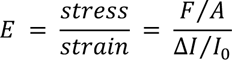

Where *E* is the Young’s modulus, *F* is the force applied on the hydrogel, *A* is the area onto which the pressure is applied, *ΔI* is the displacement, and *I_0_* is the initial thickness of the hydrogel. For all other experiments, AlgMA was dissolved at 2% (w/v) in culture medium containing 10 mmol/L of LAP, then cured with a 405 nm lamp (energy: 10 mJ/mm²).

### 2.4 Evaluation of the stability of AlgMA hydrogels

A solution of 2% [w/v] AlgMA and LAP (10 mmol/L) prepared in culture medium was cast into a mold (1.5 mm (*h)* x 5 mm (*Ø)*) and cured with a 405 nm lamp (energy: 10 mJ/mm²). The hydrogels were weighed and then incubated at 37 °C in complete culture medium. At different time points (1, 3, 14, 28 days), hydrogels were weighted after removal of the medium. The stability was determined at each time point by calculating the ratio of the hydrogel mass to its initial mass.

### 2.5 Fabrication of meltelectrowritten layers of polycaprolactone fibers

The layer of PCL fibers was designed using the Shaper software (RegenHU, Version 1.6.3) and then loaded to the HMI software for printing. The cartridge containing PCL was heated to 90 °C for 30 min to allow the polymer to melt. Rectangle-shaped perimeters of 4.6 cm (*L*, length) x 2 mm (*h*, height) were printed on glass slides in a clean, sterile environment using MEW equipment (RegenHU, R-GEN 200) through a 24G nozzle. The system’s pressure was adjusted to 110 kPa, the printing speed was 15 mm/s, the voltage was 5 kV, and the offset was 2 mm. Printed rectangles were cut into small pieces of 1 cm (*L*) and stored under sterile conditions at room temperature until their use.

### 2.6 Polycaprolactone scaffold fabrication

The PCL scaffold was designed using the Shaper software based on a histological section of a sheep lumbar IVD. Each element of the IVD (shape, location of the NP, the AF, and the various lamellae) was reproduced to be as close as possible to the structure of a native IVD using Fusion 360 (Autodesk, Version 2.0). The design was then scaled down to adapt to laboratory culture conditions and fit in 24-well culture plates using Shaper software. As a result, the PCL scaffold consisted of two components: the support and the circled PCL layers. The support was a grid of two layers of PCL fibers. The PCL cartridge was connected to a 24G nozzle and heated to 90 °C for 30 min to allow the polymer to melt. The system’s pressure was adjusted to 60 kPa, printing speed to 120 mm/s, voltage to 5 kV, and offset to 2 mm from the glass slide. Then, 13 concentric rings of 50 PCL fibers reproducing the AF part were printed on the grid. Each ring was printed using a pressure of 110 kPa, a printing speed of 30 mm/s, a voltage of 5 kV, and an offset of 1.5 mm. Once printed, PCL scaffolds were stored under sterile conditions at room temperature until use. Their dimensions (antero-posterior IVD size, transversal IVD size, antero-posterior NP size, transversal NP size, AF thickness) were measured on images taken using a digital microscope (Keyence, VR-5000) on days 0, 7, 14, and 21 to evaluate the stability of the PCL scaffold when stored at room temperature.

### 2.7 Cells isolation and culture

NP and AF cells were isolated from the lumbar IVD of a six-month-old sheep under sterile conditions (ONIRIS Veterinary School, Nantes). The separated lumbar vertebrae were thoroughly rinsed with PBS solution containing 2% (v/v) P/S, followed by removal of muscle, ligament, and other tissues to expose the IVDs. They were cut to separate AF and NP. The small pieces obtained were digested enzymatically. The tissue fragments were first incubated in HBSS containing 0.05% (w/v) hyaluronidase for 15 min at 37 °C, then in HBSS containing 0.2% (w/v) trypsin for 30 min at 37 °C and finally in culture medium containing 0.25% (w/v) collagenase for 15 h at 37 °C. The recovered suspensions were filtered through a 70-μm-pore filter and centrifuged for 5 min. The cells were counted and seeded in T75 flasks at a density of 5.10^3^ cells/cm^2^ for expansion in culture medium. Culture medium was composed of DMEM containing 10% (v/v) FBS, 1% (v/v) P/S, and 0.1% (v/v) amphotericin B. The medium was changed every 2-3 days, and cells were used or passaged until cell confluence reached 70%-80%. The cells before the fifth passage were used in the subsequent experiments.

### 2.8 Metabolic activity of AF cells on the layer of polycaprolactone fibers

Layers of PCL fibers were placed into 48-well plates. Then, 50,000 AF cells were seeded per fibrous layer, and complete culture medium was added. AF cells seeded on plastic were used as a control. All plates were maintained in a humidified incubator with 5% CO₂ at 37 °C. The medium was changed every 2-3 days. The metabolic activity of the seeded cells was assessed using an MTT assay on days 1, 7, 14, 21, and 28 following the manufacturer’s instructions. A PCL layer without AF cells was used as a negative control, and DMSO was used as a blank. Optical density values from samples were adjusted according to the blank.

### 2.9 Cell bioprinting on the polycaprolactone scaffold to obtain IVD construct

IVD cells encapsulated in AlgMA/LAP solution were deposited on PCL scaffolds using microextrusion bioprinting to produce IVD constructs. AF or NP cells were mixed with 2% (w/v) AlgMA and LAP (10 mmol/L) at a concentration of 4 x 10^6^ cells/mL. The bioinks were then transferred into a cartridge connected to a 25G needle and printed at room temperature on the IVD PCL scaffold (printing speed: 3 mm/s, pressure: 20 kPa, needle offset: 1.5 mm). The bioink with AF cells was bioprinted first between the PCL rings and cured with a 405 nm lamp (10 mJ/mm²). Then, the bioink with NP cells was bioprinted in the center of the IVD PCL scaffold and cured under the same conditions. Constructs were cultured in complete culture medium and maintained in a humidified incubator with 5% CO₂ at 37 °C. The complete culture medium is composed of DMEM containing 10% (v/v) FBS, 1% (v/v) P/S, 0.1% (v/v) amphotericin B, and 0.17 mmol/L 2AA. The medium was changed every 2-3 days. The IVD construct dimensions were measured on images taken with a macroscope on days 0, 7, 14, and 21 to evaluate their stability over time in culture.

### 2.10 Cell viability and distribution in AlgMA hydrogel

Cell viability in the IVD constructs was evaluated using a live/dead assay following the manufacturer’s instructions. Briefly, at days 1, 7, 14, 21 and 28, IVD constructs were incubated in complete culture medium containing 2 µmol/L of Calcein-AM, 4 µmol/L of Ethidium homodimer-1, and 1 µmol/L of TO-PRO^TM^-3 for 1 h at 37 °C. The medium was replaced by fresh medium, and Z-stack acquisitions of constructs were obtained using a confocal microscope (Nikon A1 R LFOV, Nikon, Champigny/Marne, France). The images were processed with Fiji software (version 2.15.1) using the “Measure” plugin. Calcein-positive NP cells were counted in the entire Z-stack. NP cell viability was calculated using the total number of nuclei in the construct and expressed as a percentage. In the meantime, NP cell sedimentation in AlgMA was also assessed. Images at 20%, 50%, and 80% of the total Z-stack height were selected to represent the bottom, middle, and top of the constructs, respectively. The number of NP cells in each image was measured and reported to the total number of NP cells counted in the three images. NP cell sedimentation was expressed as a percentage. The number of AF nuclei in the entire Z-stack was counted, and AF cell density was expressed as a number of cells/mm^3^.

### 2.11 Cell number on the polycaprolactone fibers of IVD construct

The number of AF cells on the PCL fibers in IVD constructs was evaluated using nuclear staining. Briefly, at days 1, 7, 14, 21, and 28, IVD constructs were fixed with 4% (v/v) PFA in PBS for 30 min at room temperature and labeled with Syto 13™ (1:3000) in PBS for 1 h while protected from light. Alginate lyase (10 U/mL in PBS) was added for 30 min at 37 °C under rotation to remove AlgMA. The IVD scaffolds were centrifuged at 12,000 *g* for 5 min, cut into small pieces, and transferred onto a microscope slide. Images were captured using a confocal microscope, and the number of nuclei was counted using Fiji software. AF cells on PCL fibers were expressed as a number of cells/mm².

### 2.12 Quantification of cell alignment and elongation

The morphology of cells on the layers of PCL fibers and in IVD construct were evaluated on days 1, 7, 14, 21, and 28. After removing the culture medium and rinsing with PBS, scaffolds were fixed with 4% (v/v) paraformaldehyde (PFA) in PBS for 30 min at room temperature. The cells were then permeabilized with 0.1% (v/v) Triton™ X-100 in PBS for 20 min at room temperature and stained with phalloidin-Alexa Fluor™ 647 (1:300) and Syto 13™ (1:3000) in PBS for 1 h while protected from light. For evaluating AF cells on PCL fibers in the IVD construct, the AlgMA between circles was removed using alginate lyase, as described above. Samples were observed under a confocal microscope.

Cells in 2D culture were used as a reference. Images were analyzed using Fiji software. The “analyze particles” function was used to measure cell elongation or roundness. Results are presented as scores ranging from 0 (elongated) to 1 (spherical). The “directionality” plugin (Version 2.3.0) was used to analyze cell alignment on PCL fibers. Results are expressed as a frequency of alignment. The dispersion (°) describes the standard deviation of the Gaussian, and the alignment score describes the tendance of fibers to be aligned following one direction, ranging from 0: no typical direction to 1: well oriented in one direction.

### 2.13 Scanning electron microscopy (SEM) of cells on PCL fibers and native tissues

Layers of PCL fibers at days 0, 7, and 28 cultured with AF cells, IVD construct at day 28, and native ovine AF were fixed with 4% (v/v) PFA in PBS, followed by a second fixation with 2.5% (v/v) glutaraldehyde in 0.1 mol/L in PBS for 1 h. Then, samples were incubated in 2% (v/v) osmium for 1.5 h. Then, they were washed three times with distilled water, dehydrated in an ethanol bath with increasing concentrations, and dried using short washes with different ratios of ethanol/freon. Finally, a metallization step was performed, and samples were observed using a scanning electron microscope (SEM, Gemini 300, ZEISS) at 2 kV with a 30 µm diaphragm, using the SE2 detector positioned in the chamber. Layers of PCL fibers and IVD construct images were then artificially colored using Adobe Photoshop software (Adobe Inc., 2019). The alignment of PCL fibers in layers of PCL fibers and collagen fibers in AF ovine tissues was compared. The Directionality plugin was used to analyze the angular distribution of the PCL fibers.

### 2.14 Cryogenic scanning electron microscopy (Cryo-SEM) of native ovine tissues

Samples were fixed with 4% (v/v) PFA in PBS, followed by incubation with a solution of 5% (w/v) sucrose in water for 2 h at room temperature, then 10% (w/v) sucrose in water overnight at +4 °C, and finally 15% (w/v) sucrose for 1 h at room temperature. Sucrose acts as a cryoprotectant to preserve sample structure. Samples were placed between two rivets on a sample holder attached to a shuttle. The freezing step was performed in slush nitrogen, cooled by a pressure drop (9 x 10^-2^ mBar) to −206 °C. After introduction into the preparation chamber (−140 °C), the samples were cryo-fractured by separating the two rivets, sublimated (4 min at −100 °C), and sputtered with palladium (40 s, 10 µA). The samples were then introduced into the electron microscope and positioned on the cold stage (−140 °C). Observations were performed as described above.

### 2.15 Extracellular matrix synthesis

Immunostaining was performed to assess the expression of extracellular matrix components such as type I and type II collagen by the AF cells seeded on the layer of PCL fibers and AF and NP cells in the IVD construct. On day 28, samples were fixed with 4% (v/v) PFA in PBS for 30 min at room temperature. Samples were then permeabilized and blocked by incubating with PBS containing 0.5% (v/v) Triton™ X-100 and 3% (w/v) BSA for 30 min at room temperature. Scaffolds were incubated overnight at 4 °C with primary antibodies: anti-collagen type I (1:250), anti-collagen type II (1:250). Antibodies were omitted in controls. After incubation, samples were washed with PBS containing 0.5% (v/v) Triton™ X-100 and 3% (w/v) BSA for 30 min at room temperature. Samples were incubated with anti-rabbit Alexa Fluor 568 (1:500) for 1 h at room temperature, protected from light. All samples were stained with Syto13™ (1:3000) and phalloidin-Alexa Fluor™ 647 (1:300) for 1 h at room temperature, protected from light. A final wash with PBS containing 0.5% (v/v) Triton™ X-100, 3% (w/v) BSA, and 0.02% (v/v) Tween® 20 was carried out for 30 min at room temperature, protected from light. Images were acquired using a confocal microscope, and a series of Z-projection images were reconstructed to visualize the location of the different collagen types.

### 2.16 Statistical analysis

All data were analyzed using the statistical software GraphPad Prism (GraphPad Software Inc., Version 8.0.1.). The statistical assumptions of normality and homoscedasticity were evaluated using the corresponding Shapiro-Wilk and Levene tests. A confidence level of 95% (α = 0.05) was set for all the statistical analyses. Data were analyzed with a one-way ANOVA, followed by Tukey’s post hoc test. Differences were considered significant with p-value < 0.05. The results were expressed as mean ± Standard Deviation (SD).

## 3. Results

### 3.1 Design of AlgMA biomaterial ink

Alginate-methacrylamide was used as a photocrosslinkable biomaterial ink to encapsulate the IVD cells during bioprinting. Alginate was modified with methacrylamide groups by an amidation process under aqueous conditions using DMTMM as an activating agent (**Figure 1A**). Using elemental analysis, we characterized the synthesized AlgMA, which had a degree of substitution of 20.5% ± 4.9% (N = 12, n = 2). The conditions for using AlgMA as a bioink were optimized. Two photoinitiators, LAP and Irgacure (IRG), were tested for photocrosslinking AlgMA to select the one that would produce a hydrogel with the shortest gelation time and limit cell exposure to light. LAP can effectively absorb light at 365 nm and 405 nm, while IRG only absorbs light at 365 nm [24]. A 5-second-increment photo-crosslinking experiment was used to determine the gelation time needed to form AlgMA hydrogels that can be uncasted from the mold with a spatula. Using a 2% (w/v) solution of AlgMA, we observed that LAP irradiated at 365 nm or 405 nm led to hydrogel formation in 10 and 5 s, respectively (**Figure 1B**).

**Figure 1.**
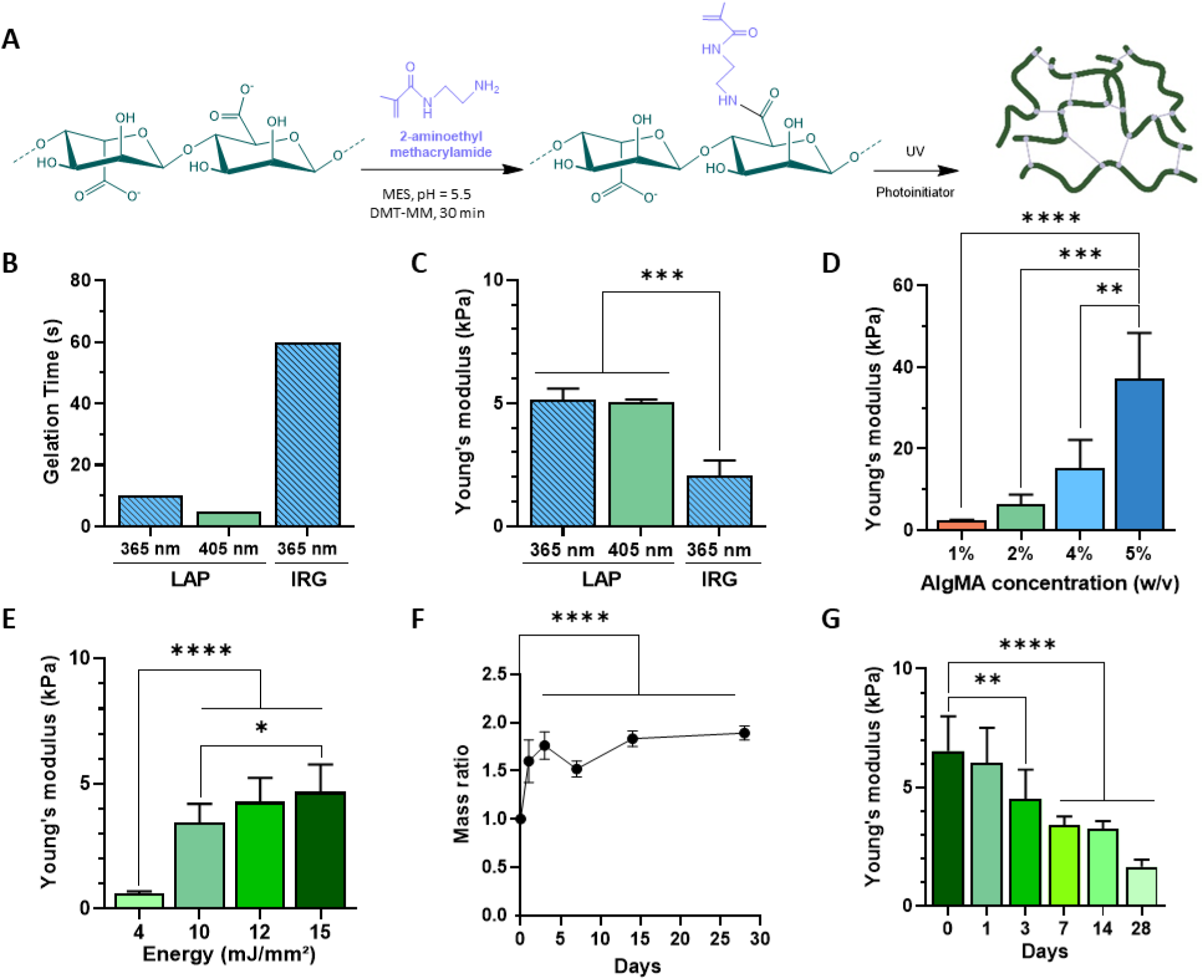
Optimization of alginate-methacrylamide (AlgMA) hydrogel for cell encapsulation. (A) Photo-crosslinkable AlgMA synthesized via amidation process. (B) Gelation time as a function of photoinitiator and irradiation wavelengths. Young’s Modulus of AlgMA as a function of (C) photoinitator (2% AlgMa, N = 1,n = 3), (D) AlgMA concentration (10 mMol/L LAP, 10 mJ/mm^2^, N = 1, n = 4), or (E) energy applied (2% AlgMa, 10 mMol/L LAP, N = 1, n = 3). (F) Swelling as a function of time and (G) associated Young’s modulus (2% AlgMAs, 10 mMol/L LAP, 10 mJ/mm2, N = 1, n = 6). Results are expressed as means ± SD. *p<0.05, **p<0.01, ***p<0.001, ****p<0.0001 indicate a significant difference.

When IRG was used as a photoinitiator, the gelation time under 365 nm irradiation was estimated to be 60 s. Together, LAP irradiated at 405 nm resulted in the fastest gelation time. The stiffness of the AlgMA hydrogels formed with the different photoinitiation conditions was evaluated using unconfined compression tests. The stiffness of the hydrogels obtained with LAP irradiated either at 365 nm or 405 nm were similar, with Young’s moduli of 5.13 kPa ± 0.46 kPa and 5.03 kPa ± 0.12 kPa, respectively (**Figure 1C**). The Young’s modulus of AlgMA hydrogels obtained with IRG was significantly lower than those obtained with LAP, with a value of 2.07 kPa ± 0.61 kPa.

Based on these results, we selected LAP irradiated at 405 nm as the photoinitiation conditions for the subsequent experiments. The influence of the AlgMA concentration and light energy (expressed as light intensity per mm^2^) on Young’s modulus was then investigated. The Young’s modulus significantly increased with the AlgMA (**Figure 1D**) concentration or the light energy (**Figure 1E**). As we were looking for a stiffness close to native heathy IVD, which is approximately 5 kPa, we decided to use an AlgMA concentration of 2% (w/v) photocrosslinked with 10 mJ/mm^2^ as the stiffness has been evaluated to be 6.25 kPa ± 2.49 kPa. Further analyses were performed to characterize the stability and mechanical properties of the photocrosslinked hydrogel. The data showed a tendency for the hydrogel to swell between day 0 and day 1 before reaching equilibrium and being stable for at least 28 days (**Figure 1F**). The swelling was accompanied by a constant decrease in Young’s modulus over the 28-day experiment (**Figure 1G**).

### 3.2 A layer of PCL fibers mimicking a single AF lamella

Native AF lamellae were first observed using cryogenic-scanning electron microscopy (cryo-SEM), revealing that AF was composed of a high-density fiber network. After tissue sectioning, we could highlight using SEM the presence of cells trapped in the ECM (**Figure 2.A**). We evaluated the relevance of MEW in producing PCL fibers that would recapitulate collagen fibers and mimic AF lamellae. The average diameter of the PCL fibers obtained with MEW was 29.74 µm ± 2.14 µm, which was different from native ovine fibers in AF lamellae measured at 0.19 µm ± 0.03 µm (data not shown). A construction of several layers of PCL fibers was printed using MEW. Fibers inside the construct tended to align relative to the horizontal axis, similar to what was observed for native AF lamellae fibers (**Figure S1**). Once in culture, PCL fibers remained cohesive throughout the 28 days of culture (**Figure 2B**). SEM images revealed that the surface appearance of PCL fibers was rough and conserved during culture. AF cells were seeded on the construct to evaluate this design’s ability to influence cell behavior and, thus, cellular organization. After 7 days of culture, the AF cells were distributed heterogeneously on the surface of the PCL fibers. Their cytoplasm was slightly extended without touching other cells (**Figure 2B**). After 28 days in culture, AF cells were closer to each other, and their cytoplasms also extended between the PCL fibers without completely covering the surface of the layer of PCL fibers.

**Figure 2.**
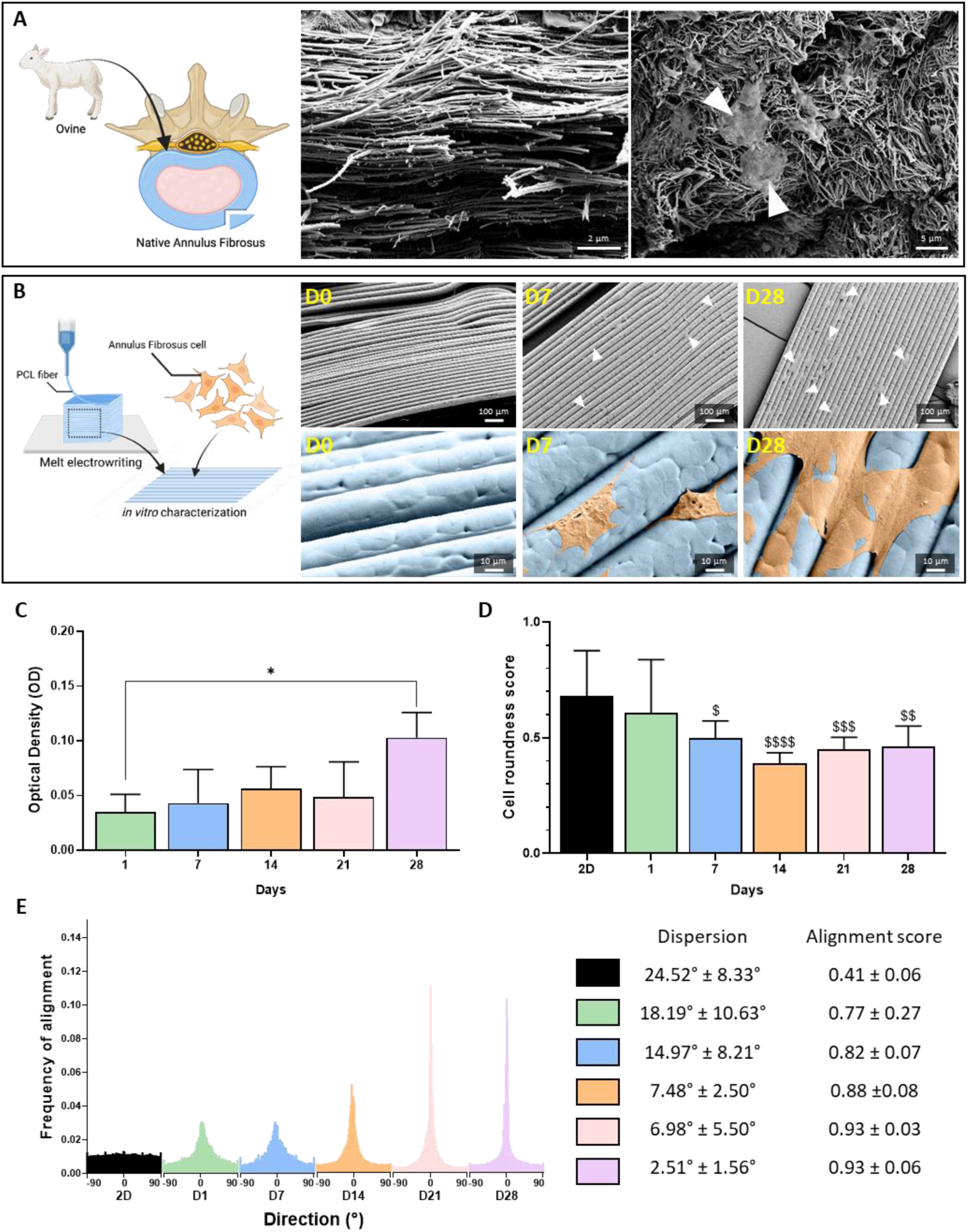
Design of polycaprolactone (PCL) layer mimicking a single ovine annulus fibrosus (AF) lamella. (A) Representative cryogenic-SEM (left) and SEM images (right) of native ovine AF tissue. (B) Representative SEM images of a PCL layer seeded with AF cells. PCL fibers: blue, cells: orange, a white head arrow indicates AF cells. (C) Metabolic activity presented as optical density (N = 1, n = 4), (D) cell roundness (N = 1, n = 5), and (E) Frequency of alignment of ovine AF cells seeded on a PCL layer as a function of time with corresponding dispersion angle and alignment score (N = 1, n = 5). Ovine AF cells cultured in 2D were used as a reference. Results are expressed as means ± SD. ^$^p<0.05, ^$$^p<0.01, ^$$$^p<0.001, ^$$$$^p<0.0001 indicate a significant difference with 2D. *p<0,05, **p<0.01 indicate a significant difference.

The metabolic activity of cells on the layer of PCL fibers was measured using an MTT assay over 28 days (**Figure 2C**). Overall, the metabolic activity of AF cells on the PCL layer tended to increase over time, and metabolic activity measured at day 28 was significantly higher than activity at day 1.

The morphology of the AF cells cultured on a PCL layer was assessed by fluorescently staining F-actin (not shown). On day 1, the cell roundness score was not different from that of AF cells seeded on a 2D plastic surface (**Figure 2D**). However, after 7 days in culture, the cell roundness score differed significantly from that of cells cultured in 2D (p<0.05). The cell roundness score decreased as the culture time increased, indicating that the cell morphology was more extended. Cells became fusiform over time, as confirmed in the SEM images (**Figure 2B**). We also confirmed the tendency for AF cells to align along the PCL fibers. On day 1, the cells did not exhibit any specific alignment direction, with an alignment score close to that of cells in 2D (**Figure 2E**). After 14 days, the AF cells were aligned along the PCL fibers with an alignment score of 0.88 ± 0.08 and a decrease in dispersion from 18.19° ± 10.63 (day 1) to 7.48° ± 2.50° (day 14). After 21 days and 28 days, the alignment score was 0.93 on both days, and the dispersion dropped from 6.98° ± 5.50° (day 21) to 2.51 ± 1.56 (day 28).

Finally, immunostaining was used to evidence the synthesis of type I and type II collagens in AF cells seeded on the PCL layer (**Figure 3**). Controls without the primary antibody showed the absence of non-specific labeling, which validated the immunostaining protocol (**Figure S2**). Intracellular labeling of type I and type II collagens was observed on all days of culture. This labeling tended to increase over culture time. These results indicate that AF cells maintained their positivity for type I and type II collagens after adhering to PCL fibers. However, there was no evidence of collagen deposition.

**Figure 3.**
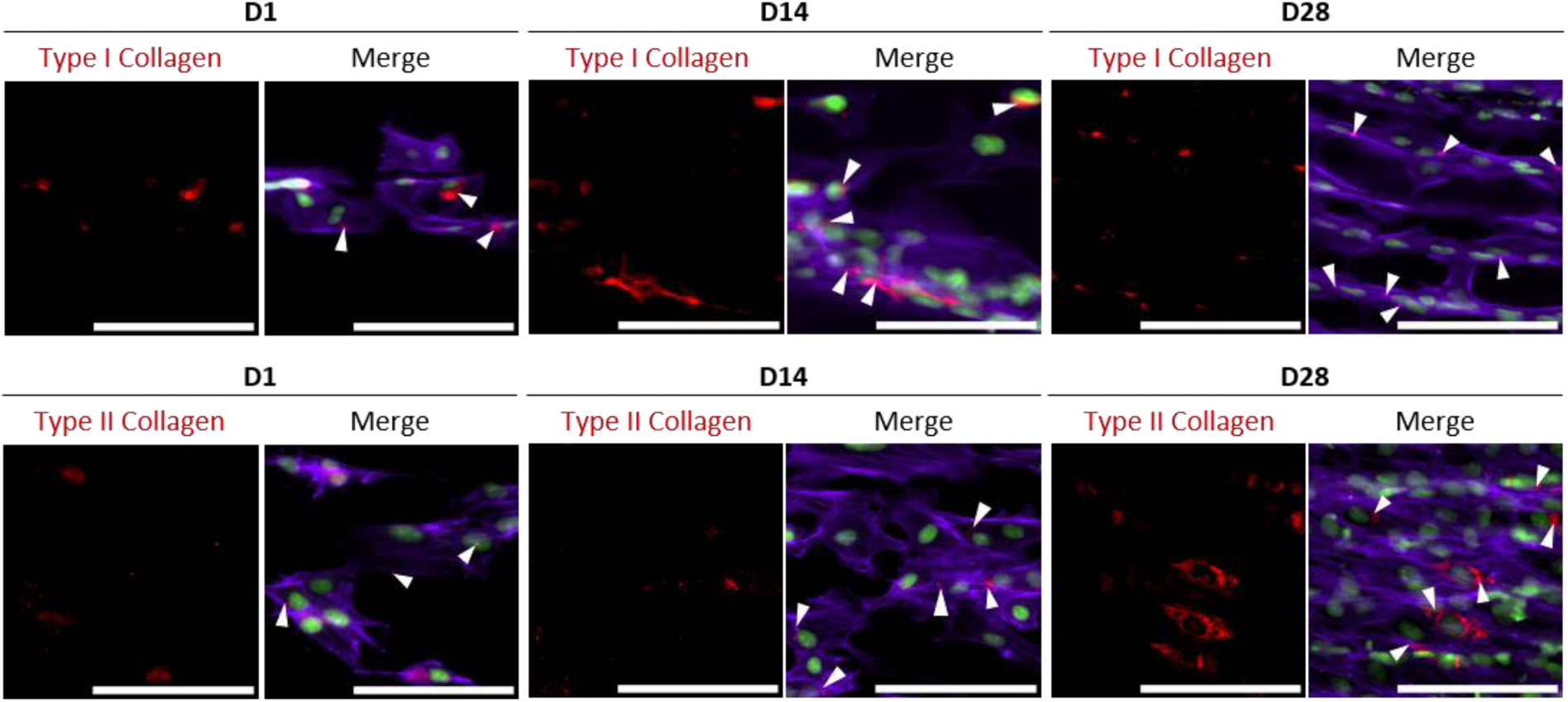
Ovine annulus fibrosus cells seeded on a polycaprolactone layer expressed type I and type II collagens. Representative Z-projection of Type I (top row) and Type II collagen (bottom row) immunostained samples at days 1, 14, and 28. Collagen: red, F-actin: magenta, nuclei: green. The white arrow indicates collagen. Scale bar: 100 µm.

Together, these findings indicate that the AF cells cultured on PCL layers underwent a morphological change from a round shape on day 1 to a fusiform shape on day 28. Moreover, cells tended to align along the PCL fibers and synthesize type I and type II collagens, suggesting the relevance of using PCL layers to guide AF cellular organization in the IVD construct.

### 3.3 A multilamellar PCL frame mimics the shape of the IVD and the lamellar organization of the AF

The MEW technique was then studied to obtain a highly resolved and reproducible frame of PCL fibers that would mimic the shape of the IVD and the lamellar organization of the AF. The PCL frame was designed to mimic the shape of the ovine IVD. The construct size was 60% of the size of an ovine IVD [25] (**Table S1**). The size was reduced to make the model suitable for culture in a 24-well culture plate. Compared with the human IVD dimensions, the construct dimension can be considered a 30% size representation (**Table S1**). Thirteen lamellae were identified in the native tissue, and each lamella was replicated using a circle of PCL. Each circle consisted of 50 PCL layers with a final theoretical height of 2 mm. Once printed, the PCL scaffold was kept at room temperature until use, and its stability was assessed (**Figure 4A**). PCL scaffolds were stable at room temperature, as there were no significant changes in their dimensions between the day of printing (day 0) and after 21 days of storage. In addition, the minor standard deviations calculated for each value indicated that the MEW technique was highly reproducible. Furthermore, the dimensions of the PCL scaffolds were not significantly different from the theoretical values set up in the software, highlighting the fidelity of the technology.

**Figure 4.**
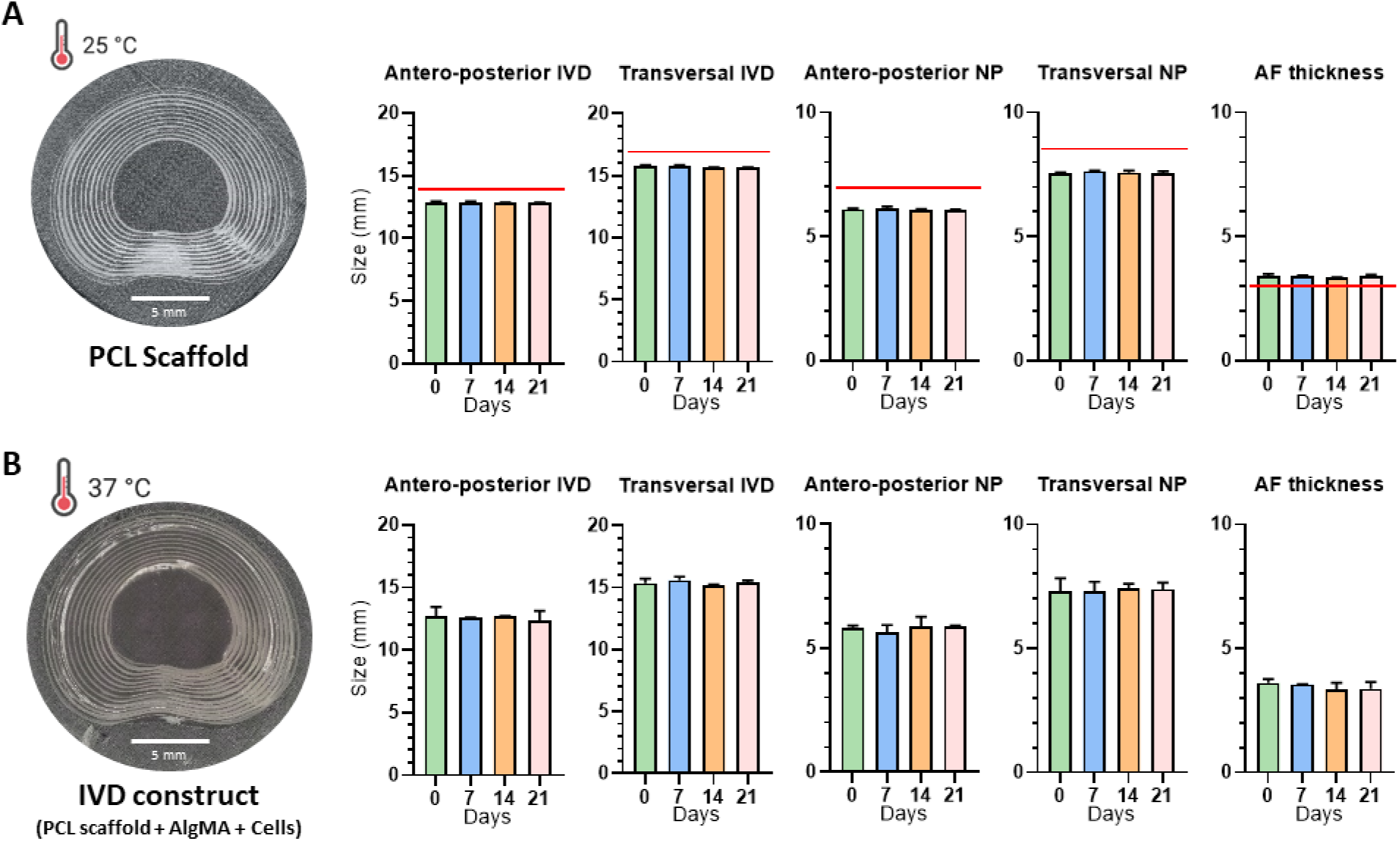
Stability of the polycaprolactone (PCL) scaffold at room temperature and of the intervertebral disc (IVD) construct in culture. Measured dimensions of (A) PCL scaffolds at 25°C (N = 1, n = 5) and (B) IVD constructs (2% AlgMA, 4×10^6^ cells/mL, N = 1, n = 2) at 37°C in culture medium as a function of time. Red bar: theoretical value. Results are expressed as means ± SD.

### 3.4 Assembly of PCL scaffold and alginate methacrylamide bioink

AlgMa bioink containing either AF (AF-bioink) or NP cells (NP-bioink) was bioprinted on the PCL scaffold to obtain the final complete IVD model. The bioprinting method enabled the control of the spatial deposition of cells and the volume of bioink deposited per sample. This minimized material loss and ensured reproducibility in construct design. The volume of each bioink deposited was estimated to be 130.10 ± 18.40 µL for the AF-bioink and 65.63 ± 2.50 µL for the NP-bioink (**Figure S3**). The stability of the cellularized IVD constructs (LAP/405 nm) was examined by measuring construct dimensions when cultured at 37 °C (**Figure 4B**). There were no significant changes in the dimensions of the IVD constructs on days 0, 7, 14, and 21, demonstrating that the IVD constructs remained stable during the culture.

#### 3.4.1 Cell viability and distribution in the IVD construct

The NP cell viability in the IVD construct was assessed with a live/dead assay. Cells bioprinted in the NP compartment of the construct remained viable for 28 days in culture, with viability close to 80% (**Figure 5A and 5B**). These results demonstrate that the bioink, bioprinting, and photocrosslinking processes are cytocompatible with NP cells. Then, NP cells at the bottom, middle, and top of the construct were quantified to evaluate cell sedimentation over culture time at 37 °C. Results showed that NP cells remained homogeneously distributed throughout the hydrogel over 14 days of culture, showing that the integrity of the polymer network was appropriated to limit cell sedimentation (**Figure 1H**). The distribution of bioprinted cells in the AF compartment was assessed by labeling the nuclei. Confocal images on day 7 showed that cell nuclei were mainly localized between PCL fibers, while after 28 days of culture, cell nuclei colocalized with PCL fibers (**Figure 5D**). These observations were confirmed by quantifying the number of AF cells in the AlgMA hydrogel and on the PCL fibers. Indeed, the number of AF cells in the AlgMA hydrogel significantly decreased between day 1 and day 21 (**Figure 5E**). In contrast, the number of AF cells on the PCL fibers increased significantly between day 1 and day 21 (**Figure 5F**). These results indicate that AF cells in AlgMA could proliferate on the PCL fibers.

**Figure 5.**
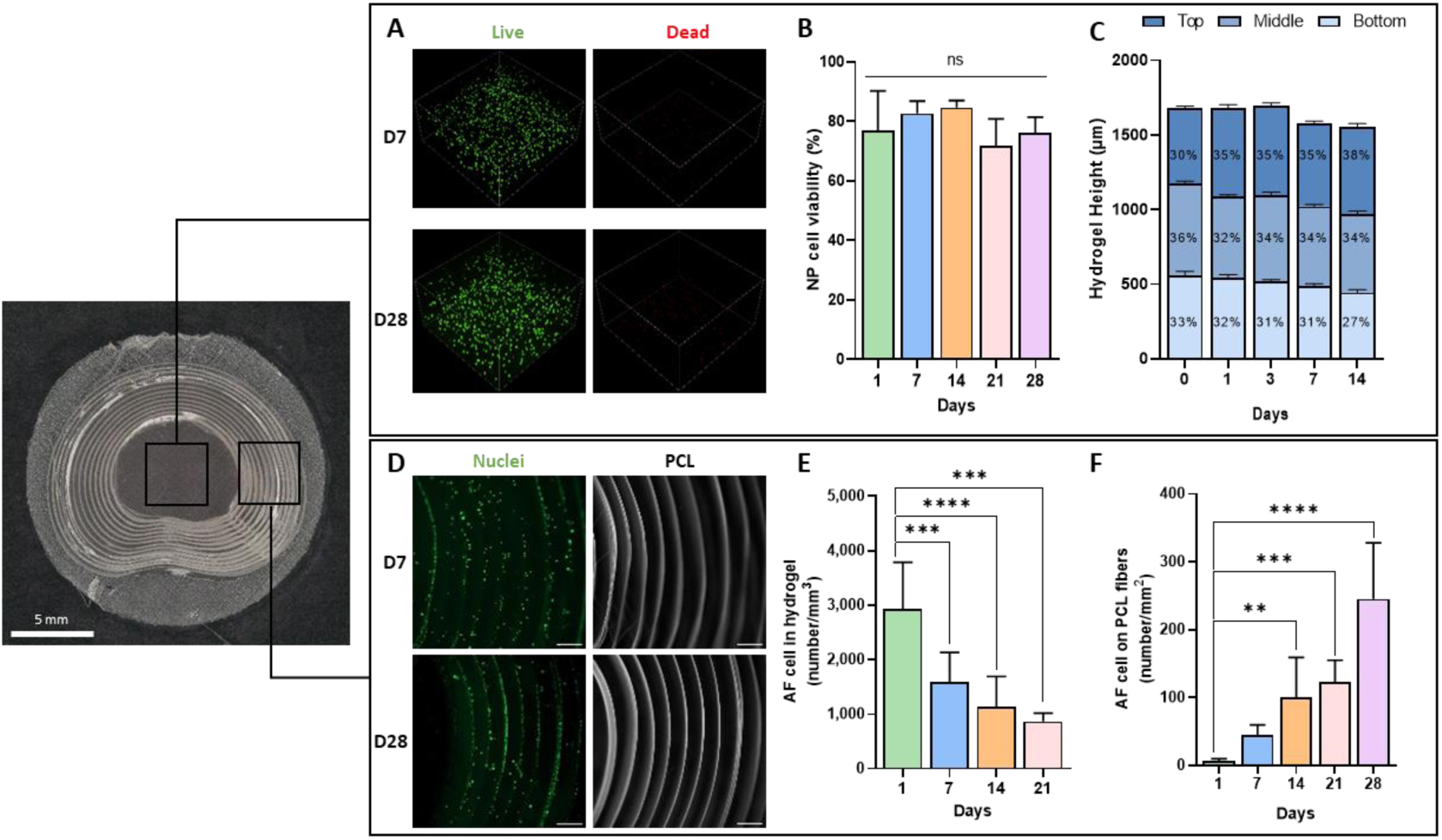
Cell behavior in the IVD construct. (A) Representative live/dead images of NP cells in AlgMA hydrogel and (B) NP cell viability (N = 1, n = 4). (C) NP cell distribution in hydrogel height (N = 1, n = 4). (D) Representative images of AF cells in the construct AlgMA hydrogel/polycaprolactone (PCL) fibers. Scale bar: 500 µm. (E) AF cell number in the hydrogel and on (F) the PCL fibers (N = 2, n = 5). Results are expressed as means ± SD. **p<0.01, ***p<0.001, ****p<0.0001 indicate a significant difference.

#### 3.4.2 Cell morphology and organization in the IVD construct

Cryo-SEM images of native ovine NP tissue demonstrated that, *in vivo*, NP cells were spherical and localized into lacunae (**Figure S5**). As we wanted to mimic native NP, we performed an SEM analysis of IVD constructs. Images revealed that, after 28 days of culture, NP cells were spherical and localized into lacunae, mimicking what was observed in native NP tissue (**Figure 6A**). The cell roundness score was quantified and calculated to be close to 1 for all culture times, meaning that NP cells were spherical (**Figure 6B**). In addition, NP cells did not exhibit any specific alignment in the hydrogel and were randomly organized in the hydrogel, mimicking their native distribution (**Figure 6C**). On the contrary, AF cells seeded in the same AlgMA hydrogel combined with the PCL scaffold exhibited a different morphology at day 28. Indeed, they displayed a fusiform shape on the PCL fibers, as shown by SEM images (**Figure 6D**).

**Figure 6.**
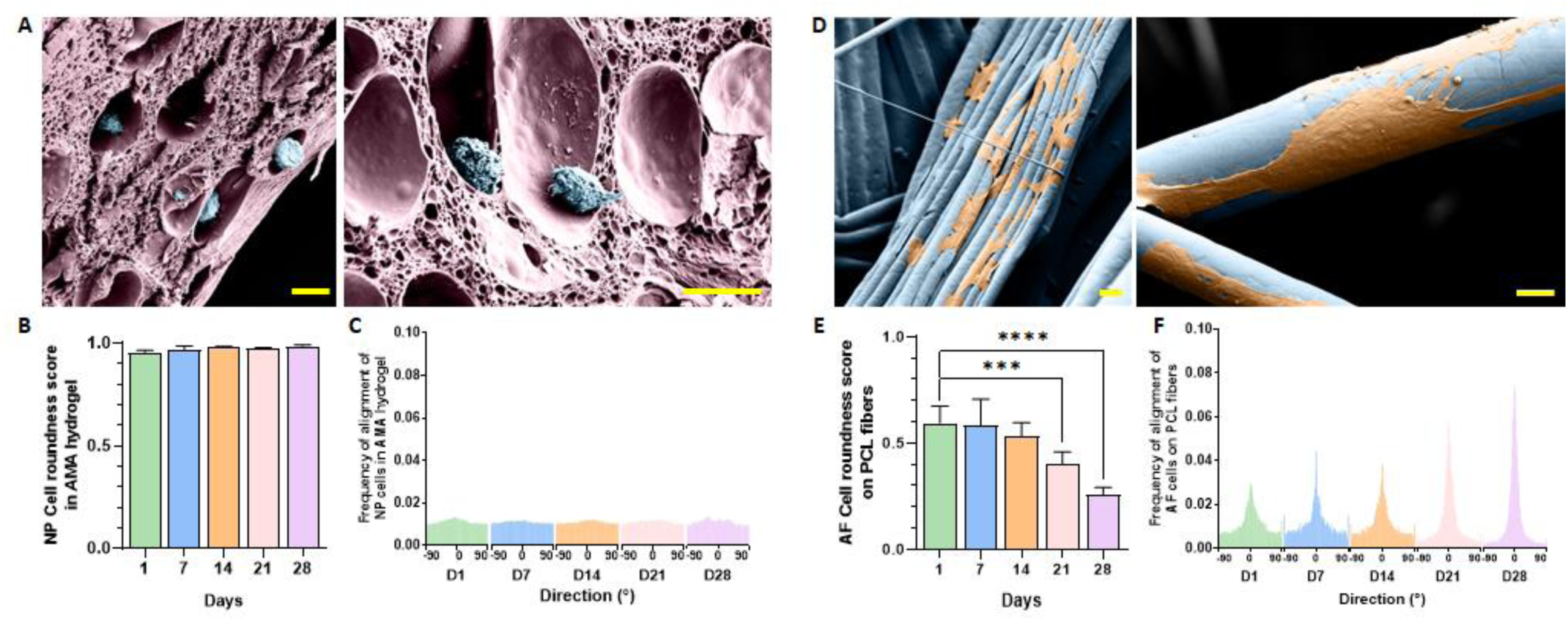
Cell morphology in the IVD construct. (A) Representative scanning electron microscopy (SEM) images of NP cells (blue) in the AlgMA (pink) hydrogel at day 28. Scale bar: 20 μm. (B) NP cell alignment and (C) roundness as a function of time (N = 1, n = 3). (D) Representative SEM images of the AF cells (orange) on the PCL fibers (blue) on day 28. Scale bar: 10 µm (left) and 20 µm (right). (E) AF cell alignment and (F) roundness on PCL fibers as a function of time (N = 1, n = 8). Results are expressed as means ± SD. ***p<0.001, ****p<0.0001 indicate a significant difference.

AF cell elongation and alignment on PCL fibers were also calculated using the same procedure as for NP cells. The cell roundness score of AF cells significantly decreased from day 1 to day 28, confirming the results observed on the layer of PCL fibers (**Figure 6E**). Moreover, the same observation was made for cell alignment. AF cell orientation along the PCL fibers increased over time, as shown by the frequency of alignment measurement (**Figure 6F**). The dispersion decreased from 14° ± 6.87° on day 1 to 6.29° ± 0.78° on day 28, while the alignment score increased from 0.83 ± 0.13 at day 1 to 0.92 ± 0.02 at day 28 (data not shown). These results confirmed that the PCL scaffold could guide AF cell orientation in the IVD construct. At day 28, immunostaining and confocal microscopy images confirmed the synthesis of type I and type II collagens by NP cells in the AlgMA hydrogel and the AF cells on the PCL fibers (**Figure 7A and 7B**), with both collagens observed within the cell cytoplasm.

**Figure 7.**
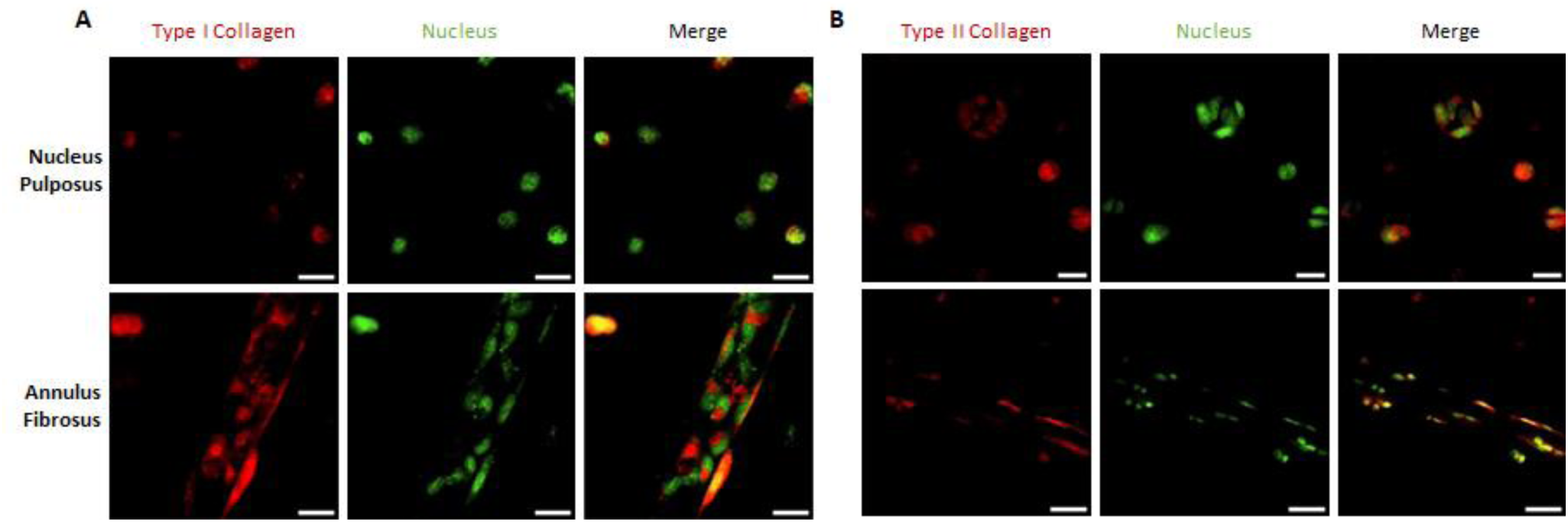
Ovine NP and AF cells seeded in the IVD construct expressed Collagen I and II after 28 days. Representative Z-projection of collagen Type I (A) and II (B) immunostained constructs at day 28. Collagen: red, nuclei: green. Scale bar: 20 µm

## 4. DISCUSSION

Currently, assessing potential therapies for disc degeneration in vitro is mainly performed on NP cells cultured in monolayer systems. Unfortunately, the reliability of these systems is limited, the 3D configuration is not reproduced, and the cells exhibit phenotypic instabilities due to the absence of an ECM-rich 3D microenvironment. Consequently, it is challenging to directly transpose the results observed on these systems to in vivo situation. The rapid development of novel treatments and the aim of reducing the use of animals in research are prompting researchers to consider potential new in vitro models that are more complex, representative and predictive. In recent years, efforts have been made to produce a 3D in vitro model by embedding cells in alginate beads [26–28]. However, these models are composed of a single cell type, and the beads’ size does not reproduce the native size of the IVD, making it impossible to use these systems to validate the delivery and efficacy of therapies. In this study, we introduced a new complete 3D IVD model, mimicking the shape of an ovine IVD and reproducing the tissue and cellular organization of the AF and NP tissues. To achieve this, we used bioprinting methods, which enable the construction of physiologically relevant structures in a controlled and reproducible manner.

Alginate was used as a polymer as it has been widely reported in the literature as a bioink [29] and has been shown to prevent IVD cell dedifferentiation [30]. Indeed, compared with other biomaterials such as chitosan, alginate promotes the accumulation of sulfated glycosaminoglycans and the deposition of type II collagen, which are essential for maintaining NP cell function [18]. However, ionically cross-linked alginate constructs lack stability and can rapidly disintegrate *in vivo* [31]. In comparison, photo-crosslinked alginate hydrogels remain stable for months [31]. In this study, we synthesized AlgMA and optimized its formulation to replicate the stiffness of the healthy NP, which was estimated to be approximately 5 kPa [32]. The mechanical properties of the hydrogel are known to influence cell behavior [17,33]. One study has shown that hydrogels with a stiffness of 20 kPa significantly induced NP cell senescence and failed to maintain the expression of their phenotypic markers [34]. In contrast, a 4 kPa hydrogel in the same study, maintained the expression of healthy phenotypic markers in IVD cells. Thus, we selected conditions where the AlgMA hydrogel exhibited a stiffness of ∼6 kPa, similar to native healthy IVD stiffness.

In our model, we used a scaffold composed of PCL fibers to mimic the lamellar organization of the native AF. PCL was employed because of its slow degradation rate, ensuring its stability throughout the culture of the *in vitro* model. A recent study on PCL electrospun fiber meshes demonstrated that in the absence of any hydrolytic or enzymatic degradation, PCL fibers remain stable for at least 90 days in PBS [35]. PCL scaffold have shown excellent cytocompatibility with disc cells and we previously explored the ability of electrospun PCL fibers to guide ovine AF cells *in vitro* and *in vivo* [36]. AF cells displayed a non-specific orientation on random fibrous PCL scaffolds, whereas PCL scaffolds with aligned PCL fibers promoted cell alignment. However, the diameter of fibers produced by electrospinning was much thinner than the fibers produced by MEW, with diameters of 1.33 µm ± 0.40 µm and 29.74 µm ± 2.14 µm, respectively. Thus, we investigated whether the AF cells would behave similarly on PCL scaffolds obtained by MEW. By seeding AF cells on a layer of PCL fibers, we demonstrated that AF cells exhibited an elongated morphology in the direction of the fibers. Moreover, the number of cells found on the fibers increased over time and cells expressed type I and type II collagens, suggesting that they maintained their differentiated phenotype. Together, these results confirmed the relevance of using layers of PCL fibers to guide AF cells in the IVD construct.

Fibrous structures mimicking that of AF are usually produced using the electrospinning technique [37–40]. However, toxic organic solvents are used, fiber deposition is random, and the formed structures are not reproducible. An alternative technique to electrospinning is melt electrowriting (MEW). Compared to electrospinning, the MEW technology allows the production of vert well-organized, high-resolution scaffolds with a defined microarchitecture using molte polymer fibers [41]. In this work, we successfully created PCL frames that precisely reproduced the shape of an ovine IVD with the different lamellae in the AF. The technique proved highly reproducible, as evidenced by the low standard deviations between different samples. It may be applicable in designing highly versatile, patient-specific models whose applications range from medical training to surgery planning and assessment of personalized therapies. It is worth mentioning that human iPSC-derived disc cells or progenitor cells could eventually be used to populate the scaffold. In addition, this technique can be combined with other bioprinting techniques, which is particularly useful for controlled cellularization of the structure. Thus, cells encapsulated in AlgMA bioink were spatially organized using bioprinting on the PCL scaffold obtained by MEW. Cells populating the construct were extracted from ovine IVDs, wherby AF and NP cells were separated. Using primary cells in the construct was encouraged by the fact that these cells represent the cell plurality of IVD and already express the tissue-specific phenotypic characteristics [42,43].

Once in culture in the construct, NP cells in the AlgMA hydrogel retained their spherical morphology and were homogeneously dispersed throughout the height of the 3D construct, exhibiting native organization. Lacunae, localized around cells, were also observed in native NP tissue and highlighted in other alginate-based hydrogels [17,44]. These studies suggested lacunae formation in the center of the hydrogels could be associated with oxygen tension or nutrient diffusion [20]. Regarding AF cells, we demonstrated that once deposited by bioprinting between the PCL rings, the number of AF cells in the hydrogel decreased as their number on the PCL fibers increased. Moreover, the cells changed their morphology from a spherical morphology in the hydrogel to an elongated morphology oriented in the direction of the PCL fibers. AF cells on PCL fibers were positive for type I and type II collagen in the construct after 28 days. The ovine cells isolated from native tissues most certainly constitute a heterogeneous population with different cellular activity or expression of collagens. Indeed, native AF comprises different zones that differ in organization, appearance, function, and matrix synthesis. The cells of the outer layers are fibroblastic and express gene markers like type I collagen. In contrast, cells of the inner layers remain rounded and possess a more chondrocyte-like phenotype, which expresses gene markers like type II collagen [45,46]. This explains why the cells in the construct are positive for these two molecules.

Although the cells in the construct were positive for type I and type II collagen markers, we could not demonstrate matrix deposition after 28 days in culture. Here, we can formulate several hypotheses. It has been suggested that the porosity of the microenvironment facilitated the diffusion of ECM molecules into the culture medium. In the absence of cells, swelling of the hydrogel was observed between day 0 and day 7, which could be linked to a low cross-linking density, as shown in studies on AlgMA hydrogels [47]. A lower cross-linking density increases the mesh size, facilitating molecule diffusion. On the other hand, current culture conditions may be insufficient to favor matrix deposition. The conditions for culturing AF cells are still not standardized and an extensive variety of culture methods exist [44]. The incorporation of native ECM components such as collagen and hyaluronic acid, and the addition of growth factors and cytokines might provide biochemical cues facilitating matrix deposition [48,49]. Nonetheless, it was shown that the culture of multi-lamellated polycarbonate urethane constructs seeded with bovine AF cells in a spinner bioreactor using media supplemented with serum, ITS, proline, dexamethasone, and pyruvate enhanced the ECM production by encapsulated AF cells in a multi-lamellae construct [40]. Applying mechanical load to the IVD construct and/or creating a localized high-density microenvironment with the macromolecular crowding technique could also enhance ECM deposition. [50]. A third hypothesis concerns the construction of the model, where overly dense PCL layers could limit the diffusion of nutrients, affecting cell functions [51]. Finally, the cells may have adopted a degenerative phenotype in our model. Studies have shown that degenerated NP cells predominantly express type II collagen intracellularly, unlike healthy cells [20]. All these hypotheses will have to be studied in future work.

The work presented here serves as a proof of concept for developing a novel 3D IVD model. Our process leverages two biofabrication methods and is fully automated and controlled, allowing for the rapid generation of a large number of models within a few hours. Existing literature has highlighted the significant impact of fiber diameter on AF stem cell morphology and gene expression [52]. For instance, AF cells exhibited nearly round morphologies on smaller-fiber-diameter scaffolds, while spindle-like cell morphologies were observed on larger-diameter scaffolds. As the fiber diameter increased, the expression levels of type I collagen in cells rose, while type II collagen and aggrecan showed opposite trends. The various parameters that affect fiber diameters are now well described, and sub-micron fibers have recently been achieved [53]. Adjusting the specific parameters in our biofabrication strategy would enable the creation of diverse models, including incorporating gradients, a feature not achievable with current tissue engineering models under development. It could be possible to reproduce the microstructural features of the inner, middle, and outer regions of native AF tissues by changing the dimensions of the fibers. Futures studies will also address the shortcomings of this work, particularly the absence of the osseous and cartilaginous endplates that are essential for ensuring the nutrition of the disc, and the induction of a degenerative state.

## 5. CONCLUSIONS

Developing a novel 3D IVD model utilizing biofabrication techniques represents a significant advancement in the field of in vitro models. By effectively mimicking the anatomical and cellular architecture of ovine IVDs, this model addresses the limitations of conventional 2D monolayer systems. We demonstrated that combining an AlgMA hydrogel and a PCL fibers scaffold can maintain the phenotype of NP and AF cells, thus fostering a more representative microenvironment for studying disc degeneration and potential therapies. Although challenges remain, such as the need for enhanced ECM deposition and optimized culture conditions, the automated and reproducible nature of this biofabrication approach opens new avenues of research and perspectives for assessing therapeutic strategies.

## Acknowledgments

The authors acknowledge the SC3M facility from the Inserm/Nantes Université/ONIRIS UMR1229 RMeS Laboratory and SFR Bonamy, Nantes, France and the IBISA MicroPICell facility (Biogenouest), member of the national infrastructure France-Bioimaging supported by the French national research agency (ANR-10-INBS-04). The authors are grateful to P. Humbert and F. Etienne for their assistance. The schematics were created using BioRender.

## Data availability

The data supporting this study’s findings are available from the corresponding author.

## SUPPLEMENTARY

**Figure S1.**
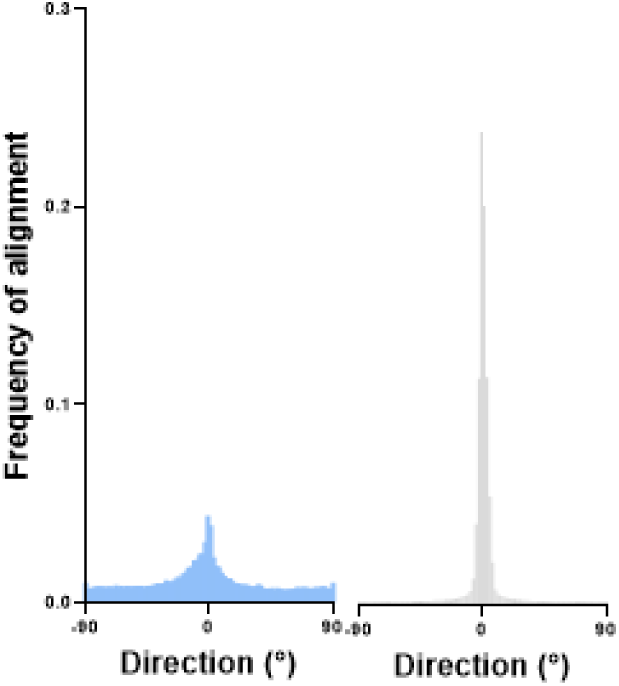
Frequency of alignment of collagen fibers in the native ovine intervertebral disc (left, N = 1, n = 2) and polycaprolatone (PCL) fibers in a PCL layer (right, N = 1, n = 44).

**Figure S2.**
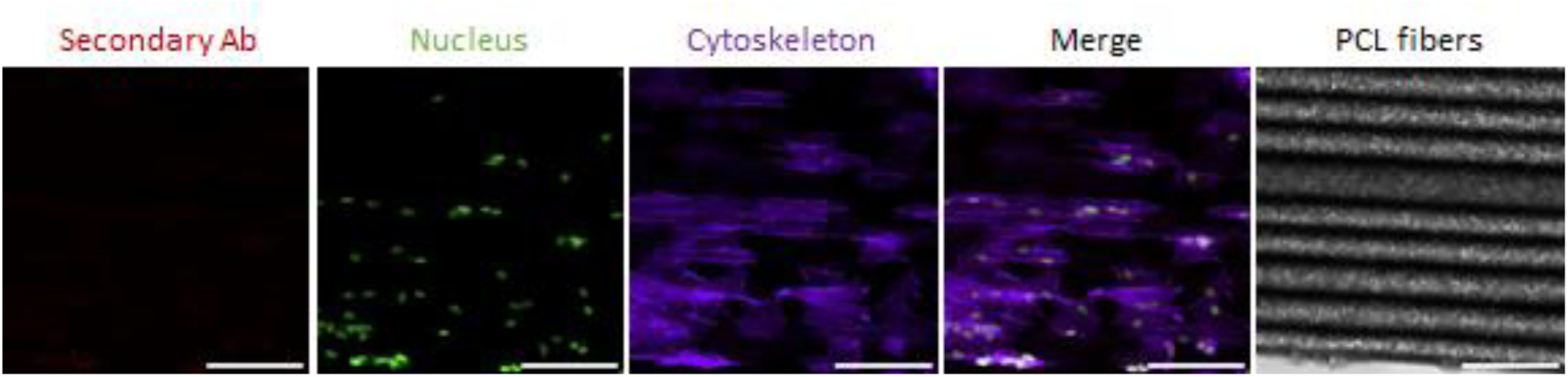
Representative images of the primary no-antibody control on ovine annulus fibrosus cells seeded on a polycaprolactone layer. Representative Z-projection of immunostained samples. Secondary antibody: red, F-actin: magenta, nuclei: green. Scale bar: 100 µm

**Figure S3.**
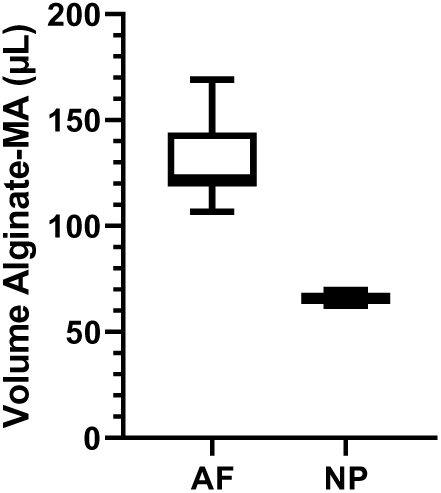
Volume of alginate-methacrylamide hydrogel deposited by extrusion in the annulus fibrosus (AF) and the nucleus pulposus (NP) regions of the intervertebral disc construct. Results are expressed as means ± SD. (N = 1, n = 11)

**Figure S4.**
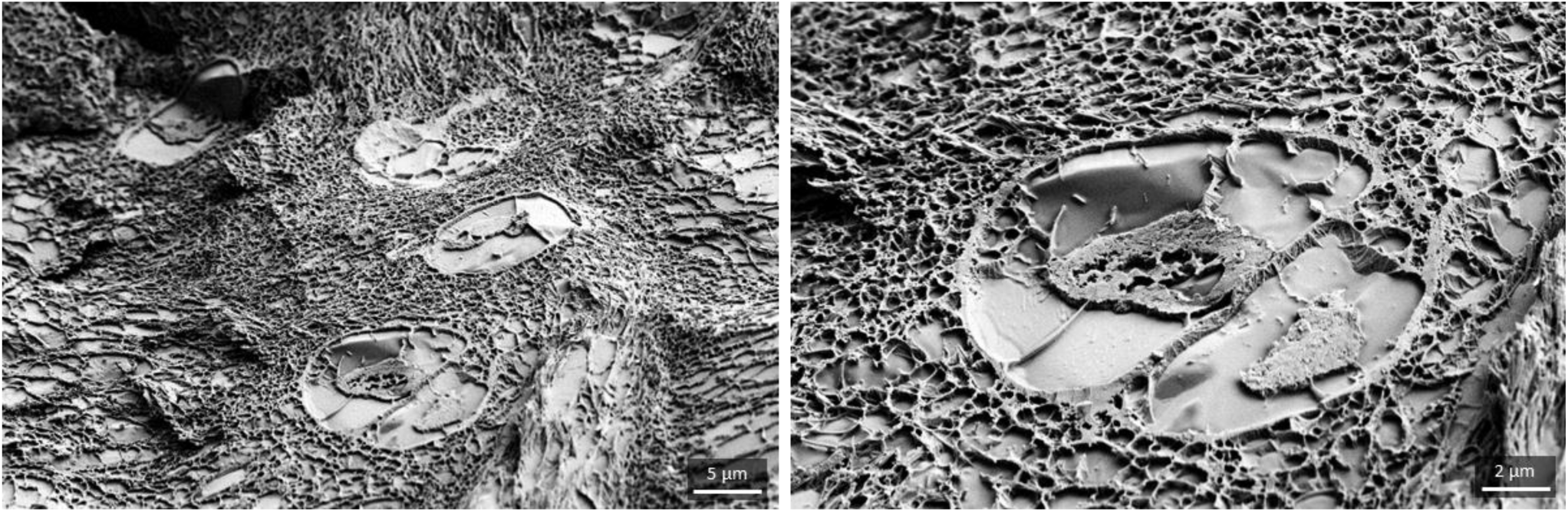
Representative cryo-scanning electron microscopy images of native ovine nucleus pulposus.

**Table S1.**
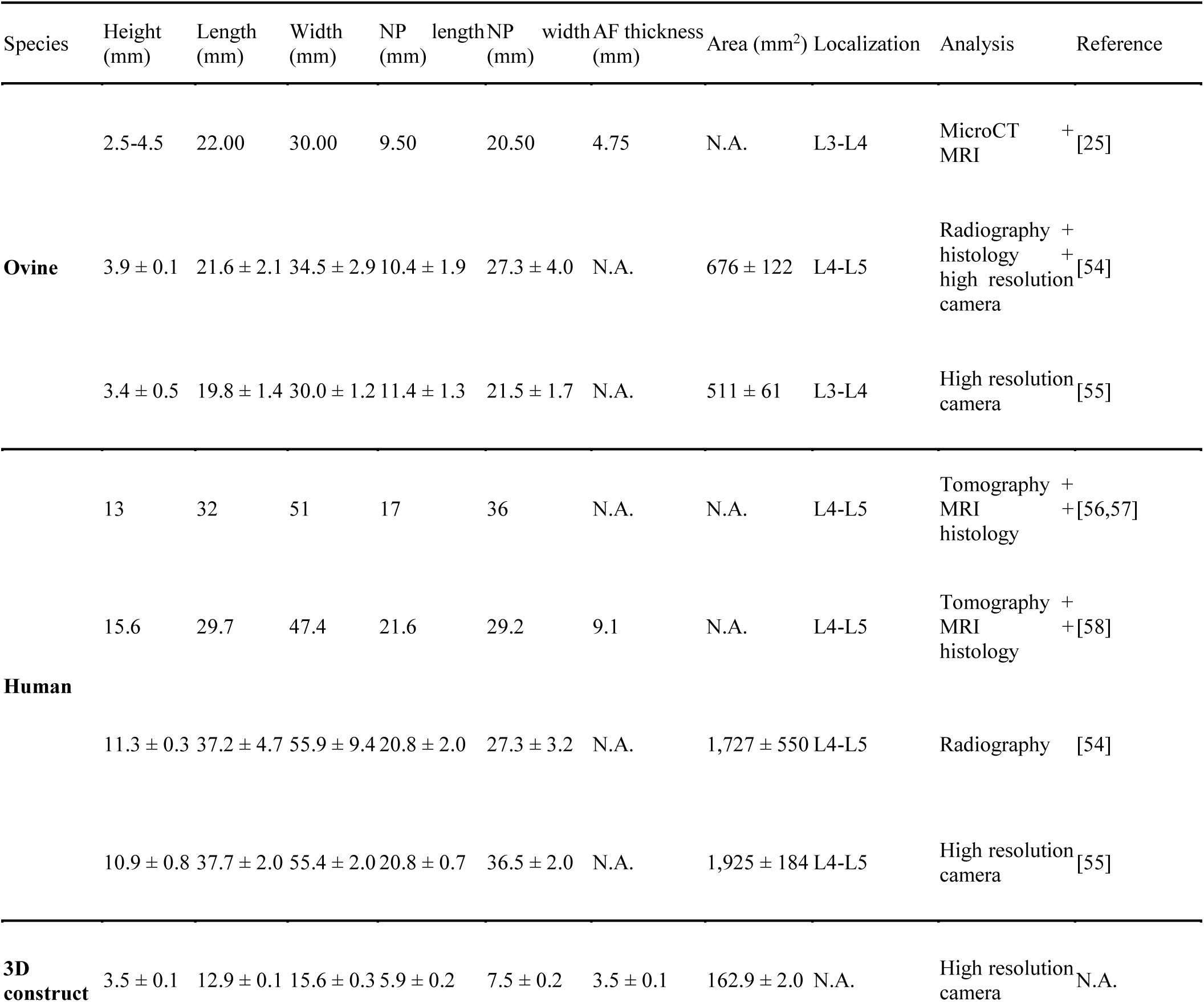
Dimensions of human, ovine, and bioprinted intervertebral discs or constructs.

## References

[1] Zhang Z, Cao K, Zhong Y, Yang J, Chen S, Li G, Wang S and Wan Z 2024 An in Vivo, Three-Dimensional (3D), Functional Centers of Rotation of the Healthy Cervical Spine World Neurosurg 184 e203–10

[2] Cloyd J M, Malhotra N R, Weng L, Chen W, Mauck R L and Elliott D M 2007 Material properties in unconfined compression of human nucleus pulposus, injectable hyaluronic acid-based hydrogels and tissue engineering scaffolds Eur Spine J 16 1892–8

[3] Schollum M L, Robertson P A and Broom N D 2009 A microstructural investigation of intervertebral disc lamellar connectivity: detailed analysis of the translamellar bridges J Anat 214 805–16

[4] Marchand F and Ahmed A M 1990 Investigation of the laminate structure of lumbar disc anulus fibrosus Spine (Phila Pa 1976) 15 402–10

[5] Costi J J, Stokes I A, Gardner-Morse M, Laible J P, Scoffone H M and Iatridis J C 2007 Direct measurement of intervertebral disc maximum shear strain in six degrees of freedom: Motions that place disc tissue at risk of injury Journal of Biomechanics 40 2457–66

[6] Liebscher T, Haefeli M, Wuertz K, Nerlich A G and Boos N 2011 Age-Related Variation in Cell Density of Human Lumbar Intervertebral Disc Spine 36 153–9

[7] Torre O M, Mroz V, Bartelstein M K, Huang A H and Iatridis J C 2019 Annulus fibrosus cell phenotypes in homeostasis and injury: implications for regenerative strategies Ann. N.Y. Acad. Sci. 1442 61–78

[8] Diwan A D and Melrose J 2023 Intervertebral disc degeneration and how it leads to low back pain JOR Spine 6 e1231

[9] Mohd Isa I L, Teoh S L, Mohd Nor N H and Mokhtar S A 2022 Discogenic Low Back Pain: Anatomy, Pathophysiology and Treatments of Intervertebral Disc Degeneration IJMS 24 208

[10] Ashinsky B, Smith H E, Mauck R L and Gullbrand S E 2021 Intervertebral disc degeneration and regeneration: a motion segment perspective Eur Cell Mater 41 370–80

[11] Cheung K M C, Karppinen J, Chan D, Ho D W H, Song Y-Q, Sham P, Cheah K S E, Leong J C Y and Luk K D K 2009 Prevalence and pattern of lumbar magnetic resonance imaging changes in a population study of one thousand forty-three individuals Spine (Phila Pa 1976) 34 934–40

[12] Fusellier M, Clouet J, Gauthier O, Tryfonidou M, Le Visage C and Guicheux J 2020 Degenerative lumbar disc disease: in vivo data support the rationale for the selection of appropriate animal models eCM 39 17–48

[13] Wu P H, Kim H S and Jang I-T 2020 Intervertebral Disc Diseases PART 2: A Review of the Current Diagnostic and Treatment Strategies for Intervertebral Disc Disease IJMS 21 2135

[14] Samanta A, Lufkin T and Kraus P 2023 Intervertebral disc degeneration—Current therapeutic options and challenges Front. Public Health 11

[15] Williams R J, Tryfonidou M A, Snuggs J W and Le Maitre C L 2021 Cell sources proposed for nucleus pulposus regeneration JOR Spine 4 e1175

[16] Kluba T, Niemeyer T, Gaissmaier C and Gründer T 2005 Human Anulus Fibrosis and Nucleus Pulposus Cells of the Intervertebral Disc: Effect of Degeneration and Culture System on Cell Phenotype Spine 30 2743–8

[17] Chou A I and Nicoll S B 2009 Characterization of photocrosslinked alginate hydrogels for nucleus pulposus cell encapsulation J Biomedical Materials Res 91A 187–94

[18] Naqvi S M and Buckley C T 2015 Differential Response of Encapsulated Nucleus Pulposus and Bone Marrow Stem Cells in Isolation and Coculture in Alginate and Chitosan Hydrogels Tissue Engineering Part A 21 288–99

[19] Shah B S, Burt K G, Jacobsen T, Fernandes T D, Alipui D O, Weber K T, Levine M, Chavan S S, Yang H, Tracey K J and Chahine N O 2019 High mobility group box-1 induces pro-inflammatory signaling in human nucleus pulposus cells via toll-like receptor 4-dependent pathway Journal Orthopaedic Research 37 220–31

[20] Sun Y, Lv M, Zhou L, Tam V, Lv F, Chan D, Wang H, Zheng Z, Cheung K M C and Leung V Y L 2015 Enrichment of committed human nucleus pulposus cells expressing chondroitin sulfate proteoglycans under alginate encapsulation Osteoarthritis and Cartilage 23 1194–203

[21] Yang S-H, Wu C-C, Shih T T-F, Chen P-Q and Lin F-H 2008 Three-dimensional Culture of Human Nucleus Pulposus Cells in Fibrin Clot: Comparisons on Cellular Proliferation and Matrix Synthesis With Cells in Alginate Artificial Organs 32 70–3

[22] Lin H A, Varma D M, Hom W W, Cruz M A, Nasser P R, Phelps R G, Iatridis J and Nicoll S B 2019 Injectable cellulose-based hydrogels as nucleus pulposus replacements: Assessment of in vitro structural stability, ex vivo herniation risk, and in vivo biocompatibility Journal of the Mechanical Behavior of Biomedical Materials 96 204–13

[23] Srivastava A, Isa I L M, Rooney P and Pandit A 2017 Bioengineered three-dimensional diseased intervertebral disc model revealed inflammatory crosstalk Biomaterials 123 127–41

[24] Lim K S, Galarraga J H, Cui X, Lindberg G C J, Burdick J A and Woodfield T B F 2020 Fundamentals and Applications of Photo-Cross-Linking in Bioprinting

[25] Casaroli G, Galbusera F, Jonas R, Schlager B, Wilke H-J and Villa T 2017 A novel finite element model of the ovine lumbar intervertebral disc with anisotropic hyperelastic material properties PLOS ONE 12 e0177088

[26] Chou A I and Nicoll S B 2009 Characterization of photocrosslinked alginate hydrogels for nucleus pulposus cell encapsulation J Biomed Mater Res A 91 187–94

[27] Sun Y, Lv M, Zhou L, Tam V, Lv F, Chan D, Wang H, Zheng Z, Cheung K M C and Leung V Y L 2015 Enrichment of committed human nucleus pulposus cells expressing chondroitin sulfate proteoglycans under alginate encapsulation Osteoarthritis and Cartilage 23 1194–203

[28] Guillaume O, Naqvi S M, Lennon K and Buckley C T 2015 Enhancing cell migration in shape-memory alginate-collagen composite scaffolds: In vitro and ex vivo assessment for intervertebral disc repair. Journal of biomaterials applications 29 1230–46

[29] Rastogi P and Kandasubramanian B 2019 Review of alginate-based hydrogel bioprinting for application in tissue engineering Biofabrication 11 #1

[30] Le Maitre C L, Freemont A J and Hoyland J A 2005 The role of interleukin-1 in the pathogenesis of human Intervertebral disc degeneration Arthritis Research & Therapy 7 R732

[31] Chou A I, Akintoye S O and Nicoll S B 2009 Photo-crosslinked alginate hydrogels support enhanced matrix accumulation by nucleus pulposus cells in vivo Osteoarthritis and Cartilage 17 1377–84

[32] Cloyd J M, Malhotra N R, Weng L, Chen W, Mauck R L and Elliott D M 2007 Material properties in unconfined compression of human nucleus pulposus, injectable hyaluronic acid-based hydrogels and tissue engineering scaffolds Eur Spine J 16 1892–8

[33] Xu P, Guan J, Chen Y, Xiao H, Yang T, Sun H, Wu N, Zhang C and Mao Y 2021 Stiffness of photocrosslinkable gelatin hydrogel influences nucleus pulposus cell properties in vitro J Cellular Molecular Medi 25 880–91

[34] Zhao R, Yang L, He S and Xia T 2022 Nucleus pulposus cell senescence is regulated by substrate stiffness and is alleviated by LOX possibly through the integrin β1-p38 MAPK signaling pathway Experimental Cell Research 417

[35] Dias J R, Sousa A, Augusto A, Bártolo P J and Granja P L 2022 Electrospun Polycaprolactone (PCL) Degradation: An In Vitro and In Vivo Study Polymers 14 3397

[36] Gluais M, Clouet J, Fusellier M, Decante C, Moraru C, Dutilleul M, Veziers J, Lesoeur J, Dumas D, Abadie J, Hamel A, Bord E, Chew S Y, Guicheux J and Le Visage C 2019 In vitro and in vivo evaluation of an electrospun-aligned microfibrous implant for Annulus fibrosus repair Biomaterials 205 81–93

[37] Martin J T, Kim D H, Milby A H, Pfeifer C G, Smith L J, Elliott D M, Smith H E and Mauck R L 2017 In vivo performance of an acellular disc-like angle ply structure (DAPS) for total disc replacement in a small animal model Journal Orthopaedic Research 35 23–31

[38] Yuan D, Chen Z, Xiang X, Deng S, Liu K, Xiao D, Deng L and Feng G 2019 The establishment and biological assessment of a whole tissue-engineered intervertebral disc with PBST fibers and a chitosan hydrogel in vitro and in vivo Journal of Biomedical Materials Research - Part B Applied Biomaterials 107 2305–16

[39] Zhu M, Tan J, Liu L, Tian J, Li L, Luo B, Zhou C and Lu L 2021 Construction of biomimetic artificial intervertebral disc scaffold via 3D printing and electrospinning Materials Science and Engineering: C 128 112310

[40] Iu J, Santerre J P and Kandel R A 2018 Towards engineering distinct multi-lamellated outer and inner annulus fibrosus tissues Journal of Orthopaedic Research 36 1346–55

[41] Liashenko I, Hrynevich A, Dalton P D, Liashenko I, Hrynevich A and Dalton P D 2020 Designing Outside the Box: Unlocking the Geometric Freedom of Melt Electrowriting using Microscale Layer Shifting

[42] Fernandes L M, Khan N M, Trochez C M, Duan M, Diaz-Hernandez M E, Presciutti S M, Gibson G and Drissi H 2020 Single-cell RNA-seq identifies unique transcriptional landscapes of human nucleus pulposus and annulus fibrosus cells Sci Rep 10 15263

[43] Riester S M, Lin Y, Wang W, Cong L, Mohamed Ali A-M, Peck S H, Smith L J, Currier B L, Clark M, Huddleston P, Krauss W, Yaszemski M J, Morrey M E, Abdel M P, Bydon M, Qu W, Larson A N, Van Wijnen A J and Nassr A 2018 RNA sequencing identifies gene regulatory networks controlling extracellular matrix synthesis in intervertebral disk tissues: RNA-Seq charactérisation of intervertebral disc J. Orthop. Res. 36 1356–69

[44] Basatvat S, Bach F C, Barcellona M N, Binch A L, Buckley C T, Bueno B, Chahine N O, Chee A, Creemers L B, Dudli S, Fearing B, Ferguson S J, Gansau J, Gantenbein B, Gawri R, Glaeser J D, Grad S, Guerrero J, Haglund L, Hernandez P A, Hoyland J A, Huang C, Iatridis J C, Illien-Junger S, Jing L, Kraus P, Laagland L T, Lang G, Leung V, Li Z, Lufkin T, Van Maanen J C, McDonnell E E, Panebianco C J, Presciutti S M, Rao S, Richardson S M, Romereim S, Schmitz T C, Schol J, Setton L, Sheyn D, Snuggs J W, Sun Y, Tan X, Tryfonidou M A, Vo N, Wang D, Williams B, Williams R, Yoon S T and Le Maitre C L 2023 Harmonization and standardization of nucleus pulposus cell extraction and culture methods JOR Spine 6 e1238

[45] Horner H A, Roberts S, Bielby R C, Menage J, Evans H and Urban J P G 2002 Cells From Different Regions of the Intervertebral Disc: Effect of Culture System on Matrix Expression and Cell Phenotype Spine 27 1018–28

[46] Bruehlmann S B, Rattner J B, Matyas J R and Duncan N A 2002 Regional BlackwellScience,Ltd variations in the cellular matrix of the annulus fibrosus of the intervertebral disc

[47] Jeon O, Bouhadir K H, Mansour J M and Alsberg E 2009 Photocrosslinked alginate hydrogels with tunable biodegradation rates and mechanical properties Biomaterials 30 2724–34

[48] Moxon S R, McMurran Z, Kibble M J, Domingos M, Gough J and Richardson S 2024 3D bioprinting of an intervertebral disc tissue analogue with a highly aligned annulus fibrosus via suspended layer additive manufacture Biofabrication

[49] Miklosic G, De Oliveira S, Schlittler M, Le Visage C, Hélary C, Ferguson S J and D’Este M 2025 Hyaluronan composite bioink preserves nucleus pulposus cell phenotype in a stiffness-dependent manner Carbohydrate Polymers 353 123277

[50] De Pieri A, Korntner S H, Capella-Monsonis H, Tsiapalis D, Kostjuk S V, Churbanov S, Timashev P, Gorelov A, Rochev Y and Zeugolis D I 2022 Macromolecular crowding transforms regenerative medicine by enabling the accelerated development of functional and truly three-dimensional cell assembled micro tissues Biomaterials 287 121674

[51] Nerurkar N L, Sen S, Huang A H, Elliott D M and Mauck R L 2010 Engineered disc-like angle-ply structures for intervertebral disc replacement Spine 35 867–73

[52] Zhou P, Wei B, Guan J, Chen Y, Zhu Y, Ye Y, Meng Y, Guan J and Mao Y 2021 Mechanical Stimulation and Diameter of Fiber Scaffolds Affect the Differentiation of Rabbit Annulus Fibrous Stem Cells Tissue Eng Regen Med 18 49–60

[53] O’Neill K L and Dalton P D 2023 A Decade of Melt Electrowriting Small Methods 7 e2201589

[54] O’Connell G D, Vresilovic E J and Elliott D M 2007 Comparison of animals used in disc research to human lumbar disc geometry Spine (Phila Pa 1976) 32 328–33

[55] Beckstein J C, Sen S, Schaer T P, Vresilovic E J and Elliott D M 2008 Comparison of animal discs used in disc research to human lumbar disc: axial compression mechanics and glycosaminoglycan content Spine (Phila Pa 1976) 33 E166-173

[56] Casaroli G, Villa T and Galbusera F 2018 Finite element comparison between the human and the ovine lumbar intervertebral disc Muscles Ligaments Tendons J 7 510–9

[57] Schmidt H, Heuer F, Drumm J, Klezl Z, Claes L and Wilke H-J 2007 Application of a calibration method provides more realistic results for a finite element model of a lumbar spinal segment Clinical Biomechanics 22 377–84

[58] Jaramillo H E, Go L and Garci J J 2015 A finite element model of the L4-L5-S1 human spine segment including the heterogeneity and anisotropy of the discs Acta of Bioengineering and Biomechanics 17 15–24

